# ARID1A-mutant and deficient bladder cancer is sensitive to EZH2 pharmacologic inhibition

**DOI:** 10.1101/2021.01.12.426383

**Authors:** James E. Ferguson, Hasibur Rehman, Darshan S. Chandrashekar, Balabhadrapatruni V. S. K. Chakravarthi, Saroj Nepal, Marie-Lisa Eich, Alyncia D. Robinson, Sumit Agarwal, Sai Akshaya Hodigere Balasubramanya, Gurudatta Naik, Upender Manne, George J Netto, Chong-xian Pan, Guru Sonpavde, Sooryanarayana Varambally

## Abstract

Metastatic urothelial carcinoma of the bladder is generally incurable by current systemic therapy. Molecular characterization of bladder cancer (BLCa) has revealed multiple candidate driver genes for BLCa tumorigenesis. Epigenetic/chromatin modifiers have been shown to be frequently mutated in BLCa, with ARID1A mutations highly prevalent in nearly 20% of early and late stage tumors. EZH2 is a histone methyltransferase that acts as an oncogene. The data herein show that ARID1A deficient tumors, but not ARID1A wild-type tumors are sensitive to EZH2 inhibition. Specifically, EZH2 inhibitor-treated ARID1A deficient bladder cancer cells show significantly reduced cell viability, colony formation, and *in vivo* tumor growth relative to ARID1A-wild type bladder cancer cells. Thus, our study suggests that a specific subset of bladder cancer patients with ARID1A mutations can be therapeutically treated with pharmacologic inhibitors targeting EZH2.

## Introduction

Bladder cancer (BLCa) is the 6th most common cancer in the US, and leads to ∼18,000 deaths annually (1). Unfortunately, bladder cancer outcomes have been relatively stagnant despite the recent introduction of a number of new therapies including immunotherapy. Next generation sequencing has revolutionized our understanding of bladder cancer and provides an opportunity to develop personalized therapy (2, 3). These analyses have revealed that epigenetic modifying genes are frequently mutated in bladder cancer as over 90% of tumors harbor inactivating mutations in at least one chromatin modifying enzyme (4). About 20% of BLCa have truncating and inactivating mutations in the AT Rich Interactive Domain 1A (ARID1A) gene, a member of the SWI/SNF chromatin modifying complex (aka BAF in mammals), making it the most frequently mutated epigenetic gene in bladder cancer. The development of such epigenetic mutations is one of the early events in BLCa tumorigenesis (5).

ARID1A is the DNA-binding component of the large 1.15 MDa SWI/SNF complex. This complex contains ATPase activity which is important for ATP-dependent chromatin remodeling that generally results in increased transcriptional accessibility and modulates diverse gene programs and cellular processes including DNA repair, telomere cohesion, and immune recognition (reviewed in refs (6, 7)). Thus, the functional ramifications of ARID1A deficiency are dependent on its downstream transcriptional consequences, which can be altered by other epigenetic transcriptional regulators and the specific cellular context.

Previously, we and others have shown that the histone methyltransferase Enhancer of Zeste Homolog 2 (EZH2), which generates a transcriptionally repressive chromatin mark, is over-expressed in many aggressive cancers where it is thought to drive growth and is thus considered an oncogene (8-11). EZH2 functions as the catalytic subunit of the polycomb repressive complex 2 (PRC2) which trimethylates lysine 27 on histone 3 (H3K27Me3), resulting in transcriptional silencing of numerous genes including tumor suppressors (12). EZH2 expression is increased in aggressive bladder cancer and promotes bladder cancer cell proliferation (13, 14).

Interestingly, work in drosophila, yeast, and ovarian clear cell carcinoma has revealed a functional antagonism between ARID1A and EZH2, and that mutations in ARID1A sensitize cells to EZH2 pharmacologic inhibition with the small molecule inhibitor GSK-126 both *in vitro* and *in vivo* (15-17). We hypothesized that bladder cancer cells with ARID1A mutations would show sensitivity to EZH2 inhibition which could be utilized as a therapeutic target in patients with ARID1A-deficient bladder cancer.

Herein, we show that ARID1A-deficient BLCa is particularly sensitive to inhibition of EZH2 with the small molecule inhibitor GSK-126. Specifically, EZH2 inhibitor-treated ARID1A mutant and/or knockdown cells show significantly reduced cancer cell viability, colony formation, and xenograft growth relative to ARID1A-wild type cells. These data support the rationale for re-purposing inhibitors of EZH2 to treat patients harboring advanced bladder cancers with ARID1A mutations.

## Materials and Methods

### Cell lines and reagents

HEK293T (ATCC) and all bladder cancer cell lines HT1197 (ATCC, Manassas, VA, USA), HT1376 (ATCC), T24 (ATCC), 5637 (ATCC), RT112, VM-CUB1 (DSMZ, Braunschweig, Germany) were grown in Dulbecco’s 90% Dulbecco’s MEM (4.5 g/L glucose) with penicillin– streptomycin (100 U/ml) and 10% fetal bovine serum (Sigma-Aldrich, St Louis, MO, USA) in 5% CO_2_ cell culture incubator. GSK-126 was obtained from Med Chem Express (catalog # HY-13470).

### Human bladder cancer lysate preparation

With IRB approval, and after pathologic stage and grade determination, protein lysates were prepared from human bladder cancer samples and surrounding normal mucosa, using samples from radical cystectomy (as previously described in ref (18)).

### *In silico* data analysis

Using cBioPortal, the mutation profiles of ARID1A and EZH2 in muscle-invasive and non-muscle invasive bladder cancer were obtained. cBioPortal provides user-friendly graphical interface to analyze whole exome sequencing datasets from The Cancer Genome Atlas Project and other published reports (2, 5, 19, 20). To study EZH2 gene expression profile in bladder invasive carcinoma patients, TCGA level 3 RNA-seq data (including “raw_read_count” and “scaled_estimate” for each sample) was downloaded for all primary tumor and normal samples using TCGA-Assembler (21). Transcript per million values for each gene was obtained by multiplying scaled estimate by 1,000,000. Using patient ID from cBioPortal, primary tumors were categorized based on ARID1A mutation status. Boxplot was generated using R (https://cran.r-project.org/).

### ARID1A shRNA

High titer lentivirus was generated by transfecting HEK293FT cells with a mixture containing three plasmids and 25-kDa linear polyethyleneimine (PEI) (Polysciences, Inc. PA, USA, 239662). In brief, 2.5 µg pMD2.G (Addgene, MA, USA, 12259), 6.5 µg psPAX2 (Addgene, 12260) and 3 µg pLKO.1-shARID1A (MISSION shRNA (Sigma-Aldrich), (TRCN0000059090 or TRCN0000059089) were diluted in 1.5 ml Opti-MEM (reduced serum) (ThermoFisher, 11058021) medium and incubated for five minutes. Afterwards, 36 µl of PEI was added to the plasmid mixture and incubated for 20 min at room temperature. The plasmid/PEI mixture was added to the HEK293FT cells (70-80% confluence) grown in a T-75 flask and incubated in 5% CO_2_ at 37°C overnight. The original medium was replaced with fresh medium [DMEM, 1x MEM Non-essential amino acid (ThermoFisher, 11140050) and Fetal Bovine Serum (ThermoFisher, 10082147) 18-20 h after transfection. The supernatant containing the first batch of the lentivirus was collected 24 h after replacement of the medium. This step was repeated and the second batch of lentivirus was collected after 48 h. The two batches of lentivirus were combined and filtered through a 0.45 μm filter. To concentrate the lentivirus, the filtrate was placed in a centrifuge tube containing Opti-prep (∼4 ml) (Sigma, D1556) at the bottom as the cushion and centrifuged at 50,000 × g for 2h using an SW32Ti rotor (Beckman Coulter). After centrifugation, a layer containing the lentiviral particles located between the medium and Opti-prep was collected and placed in a 50-ml Falcon tube. Culture medium was added to the tube to increase the volume to 50 ml. A second centrifugation was done at 5000 × g overnight at 4°C. The pellet containing the lentiviral particles was re-suspended in ice-cold PBS (pH 7.4) and stored as 10 μl aliquots at – 80°C. These lentiviruses were used to infect ARID1A wild-type BLCa cell lines (T24, 5637 and RT112) which were then selected for stably transfected clones using puromycin (Gibco, ThermoFisher), at the concentration of 1-2µg/ml for 2 weeks. Stable clones were selected and tested for ARID1A knockdown using the methods below.

### Immunoblot analyses

Antibodies used are noted in Table 1 below. All antibodies were used at dilutions optimized in our laboratory. For immunoblot analysis, protein samples were separated on a sodium dodecyl sulfate– polyacrylamide gel electrophoresis and transferred onto Immobilon-P PVDF membrane (EMD Millipore, Billerica, MA, USA). The membrane was incubated for 1 h in blocking buffer (Tris-buffered saline, 0.1% Tween (TBS-T), 5% nonfat dry milk), followed by incubation overnight at 4 °C with the primary antibody. After a wash with TBS-T, the blot was incubated with horseradish peroxidase-conjugated secondary antibody and signals were visualized by Luminata Crescendo chemiluminescence western blotting substrate as per the manufacturer’s protocol (EMD Millipore).

**Table 1:** List of antibodies used in this study, Related to the Materials and methods.

| **Antibody** | **Application** | **Dilution** | **Supplier** | **Cat. No.** |
| --- | --- | --- | --- | --- |
| **ARID1A** | **IB** | IB, 1:1000 | Cell Signaling Technology, Danvers,<br>MA | D2A8U |
| **EZH2** | **IB** | IB, 1:1000 | Cell Signaling Technology, Danvers,<br>MA | 5246S |
| **H3K27Me3** | **IB** | IB, 1:2000 | Cell Signaling Technology, Danvers,<br>MA | 9733S |
| **Histone H3** | **IB** | IB, 1:2000 | Cell Signaling Technology, Danvers,<br>MA | 4499S |
| **β-Actin** | **IB** | IB, 1:100000 | PTG lab, Rosemont, IL | HRP-60008 |

### Cell proliferation assays

Cell proliferation was measured by luminescence. For this, either ARID1A-mutant bladder cancer cells – HT1197, HT1376 and VMCUB1, or wild-type cells – T24, RT112, and 5637 were used. These cells were treated with DMSO, 7.5 or 10 µ*M* of GSK-126 for 8 days while replenishing with the fresh drug + growth medium every other day. Measurements were made according to manufacturer’s instructions. Briefly, plates were removed from the incubator and allowed to equilibrate at room temperature for 30 minutes, and equal volume of CellTiter-Glo^®^ 2.0 reagent was added directly to the wells. Plates were incubated 2 minutes on a shaker to induce cell lysis and then allow the plates to incubate at room temperature for 10 minutes to stabilize the luminescent signal. Luminescence was measured on a Synergy HTX multi-mode reader (BioTek Instruments, Inc., Winooski, VT, USA).

### Colony formation assay

Bladder cancer cells were seeded at 800 cells per well of 6-well plates (triplicate) and incubated at 37 °C with 5% CO_2_ for 7–10 days while treating with GSK-126 every other day. Here both untreated and DMSO treated cells served as controls. Colonies were fixed with 10% (v/v) ethanol for 30 min and stained with crystal violet (Sigma-Aldrich, St Louis, MO, USA) for 20 min. Then, the photographs of the colonies were taken using Amersham Imager 600RGB (GE Healthcare Life Sciences, Pittsburgh, PA, USA). Colony quantification was carried out using ImageQuant TL Colony v.8.1 software (GE Healthcare Life Sciences).

### Xenografts tumor growth assay

Animal experiments were approved by the Institutional Animal Care and Use Committee of UAB. For tumor xenograft experiments, NU/J nu/nu mice aged 6-8 weeks (n = 5 for each group) from Jackson Laboratories were injected subcutaneously into the right dorsal flanks with human bladder cancer cell lines harboring ARID1A wild-type alleles (RT112 and 5637), ARID1A mutant alleles (HT1376 and VMCUB-1), or stable ARID1A knockdown (RT112 and 5637) (1-2 × 10^6^ cells in 50 µL of incomplete media without FBS, and 50 µL of Matrigel). After inoculation of the cells, tumor growth was measured with Vernier calipers and recorded on a weekly basis. Tumor volume was calculated with formula: 0.5 × tumor length × tumor width^2^. For all *in vivo* studies, GSK-126 was administered intraperitoneally at a dose of 100mg/kg once daily. The final volume of drug/vehicle was 0.2 ml per 20 g body weight in 20% captisol adjusted to pH 4-4.5 with 1 N acetic acid. Treatment with GSK-126 for a duration of 21 days was started after the tumor volume reached to 150 to 200 mm^3^ in size. At the end of the treatment, tumors were excised, weighed, processed and stored for downstream molecular analysis.

## Results

### Dysregulation of ARID1A in bladder cancer

We and others have shown that EZH2 is critical for tumor cell survival, tumor growth, and regulates tumor suppressor gene and microRNA expression in aggressive prostate, breast, bladder and other cancers (8, 9, 22). Our *in silico* mutation analysis via cBioPortal (http://www.cbioportal.org/), using the Memorial Sloan Kettering Cancer Center bladder cancer sequencing dataset and the TCGA dataset, indicated that up to 28% of bladder cancers harbor nonsense or truncating mutations in ARID1A (**Fig. 1A**). Furthermore, using protein lysates from tumors isolated from patients, we found that EZH2 protein levels were dramatically increased in tumors compared to surrounding normal urothelial mucosa, while ARID1A showed the opposite expression pattern (**Fig. 1B**). Histone H3-trimethylated lysine 27 levels were increased in tumor samples as expected, which correlated with overexpression of EZH2 (**Fig. 1B**). Furthermore, ARID1A-mutated bladder cancers express high level of EZH2 (**Fig. 1C**).

**Figure 1:**
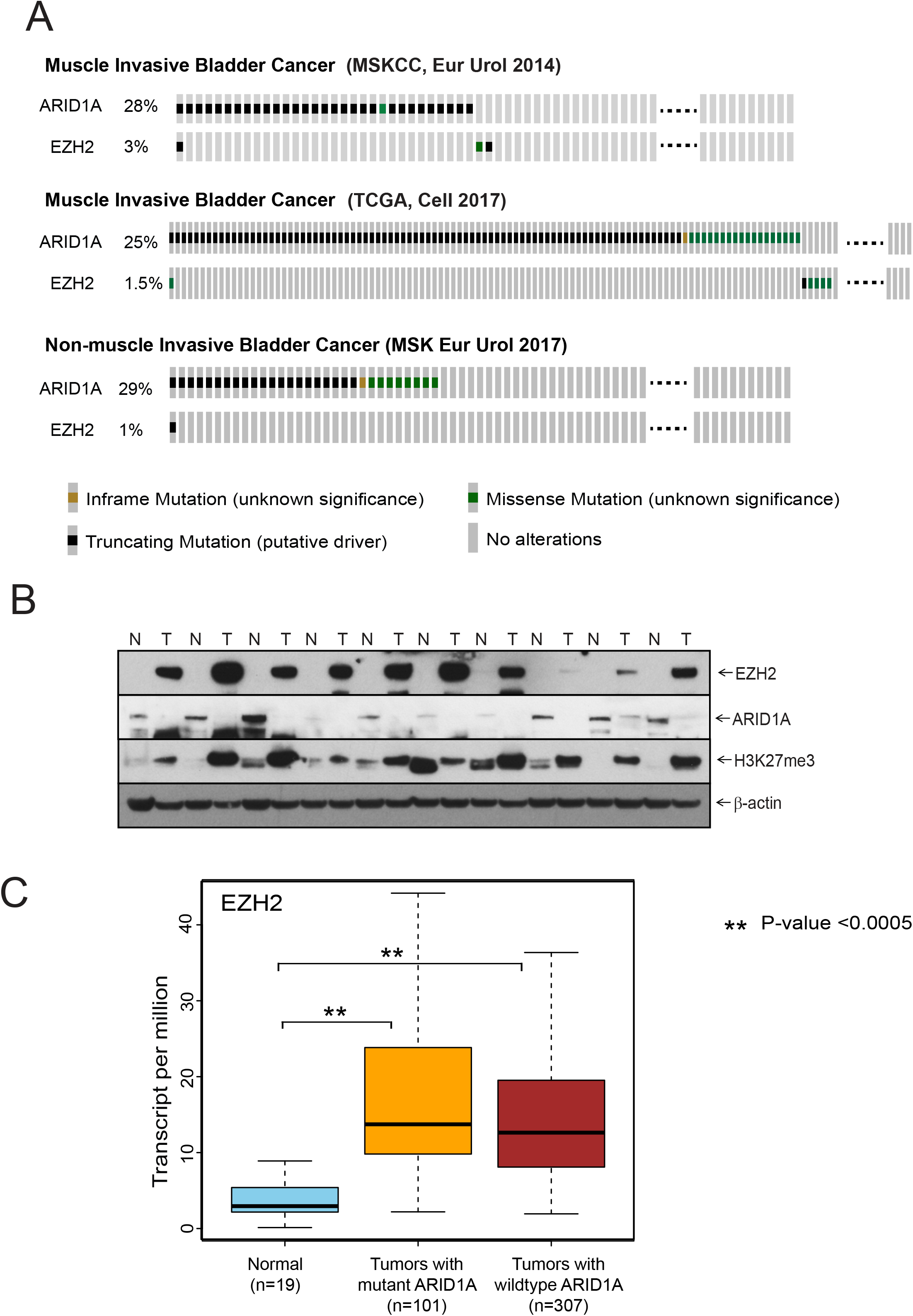
Bladder cancers harbors frequent inactivating mutations in ARID1A, and show low levels of ARID1A protein, and elevated EZH2 protein. (A) Oncoplots showing prevalence of ARID1A and EZH2 mutations in muscle invasive and non-muscle invasive bladder carcinoma. Mutation profile from three independent genomic sequencing data gathered from cBioPortal.org show significant number of bladder cancers harbor truncating or missense mutations in ARID1A. (B) Immunoblot of invasive bladder tumor (T) and matched normal tissue (N) (from cystectomy specimens) shows expression pattern of EZH2 and ARID1A. Histone H3 trimethyl lysine 27 (H3K7me3) level is increased in BLCA corresponding to overexpression of EZH2. (C) Box-whisker plot showing expression level of EZH2 in TCGA bladder carcinoma tumors with wild type and mutant ARID1A.

Our analysis of COSMIC cell line dataset (Wellcome Trust Sanger Institute, UK) suggested that multiple bladder cell lines harbored mutations in ARID1A. Among them, HT1197 has non-sense and missense substitution, HT1376 has frameshift deletion and missense substitution, and VM-CUB1 has a non-sense substitution mutation in ARID1A gene. Other bladder cancer cell lines (T24, RT-112 and 5637) do not harbor mutations in the ARID1A gene. These data have been independently confirmed by other groups (23). Importantly, these cell lines harboring truncating mutations in ARID1A show a dramatic decrease in levels of the ARID1A protein (**Fig 2A**).

**Figure 2:**
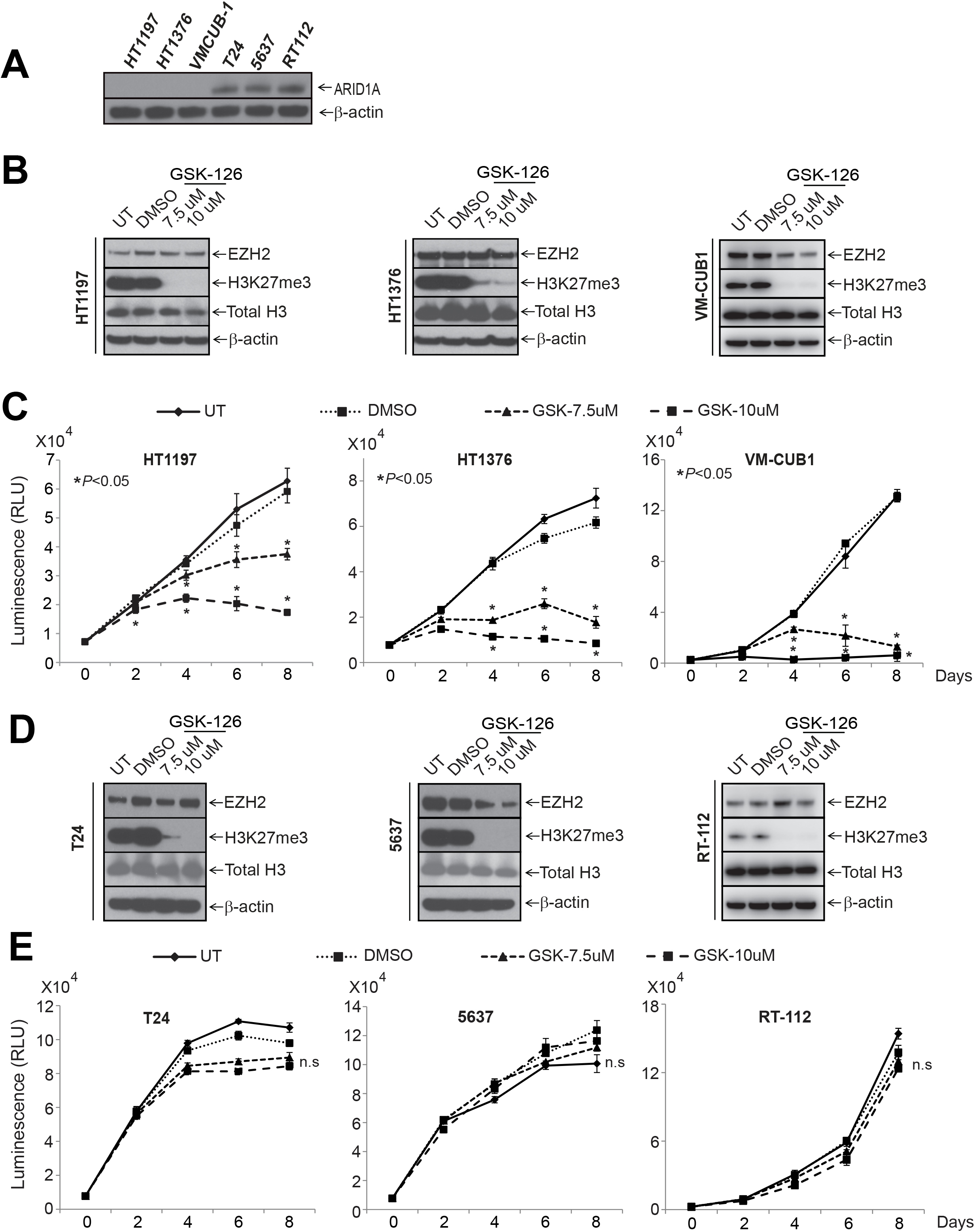
Bladder cancer cell lines with ARID1A mutation are sensitive to EZH2 inhibition. (A) Immunoblot shows expression of ARID1A in different bladder cancer cell lines including ARID1A mutant (HT1197, HT1376, and VMCUB-1), and ARID1A wild type (T24, 5637, and RT112) cell lines. (B) Immunoblot analysis showing expression level of EZH2, Histone H3 trimethyl lysine 27 (H3K27me3) and total Histone 3 in ARID1A mutant bladder cancer cell lines (HT1197, HT1376 and VM-CUB1) after treatment with EZH2 small molecule inhibitor GSK126. (C) Cell proliferation assay indicated ARID1A mutant bladder cancer cells are sensitive to GSK126. (D) Immunoblot analysis as above in (B) in ARID1A wild type bladder cancer cell lines (T24, 5637 and RT-112) after GSK126 treatment. (E) Cell proliferation assay of ARID1A wild type bladder cancer cells showed no effect of GSK126 treatment. “ns” – non-significant.

### ARID1A mutation is associated with sensitivity to EZH2 inhibitors

GSK-126 is a specific small molecule inhibitor of EZH2. To investigate the effect of GSK-126 on cell proliferation in ARID1A mutated bladder cancer cells, we performed cell viability experiments in cell lines with and without mutations in ARID1A. GSK-126 treatment significantly decreased cell viability in ARID1A mutated bladder cancer cell lines HT1197, HT1376 and VM-CUB1. However, no significant change in cell number was seen in ARID1A wild type cells (**Fig.2 C, E**). To confirm that the doses used were effective in inhibiting the histone methyltransferase activity of EZH2, we performed immunoblot analysis for the EZH2 substrate H3K27Me3. As expected, GSK-126 decreased H3K27Me3 in all bladder cancer cell lines (**Fig. 2 B, D**). Thus, at these concentrations, ARID1A mutant bladder cancer cell lines are sensitive to EZH2 inhibition with GSK-126, while ARID1A wild-type cell lines are resistant.

To investigate these findings *in vivo*, we performed mouse xenograft experiments with ARID1A mutant and wild-type bladder cancer cell lines. Indeed, while xenografts harboring ARID1A mutations were sensitive to systemic GSK-126 treatment, xenografts with wild-type ARID1A were resistant (**Fig 3 A-D, supp Fig 1 A-C**). To confirm that EZH2 was still pharmacologically active in the ARID1A wild-type xenografts, protein lysates were subjected to H3K27Me3 immunoblotting (**Supp Fig 2 A-C**)

**Figure 3:**
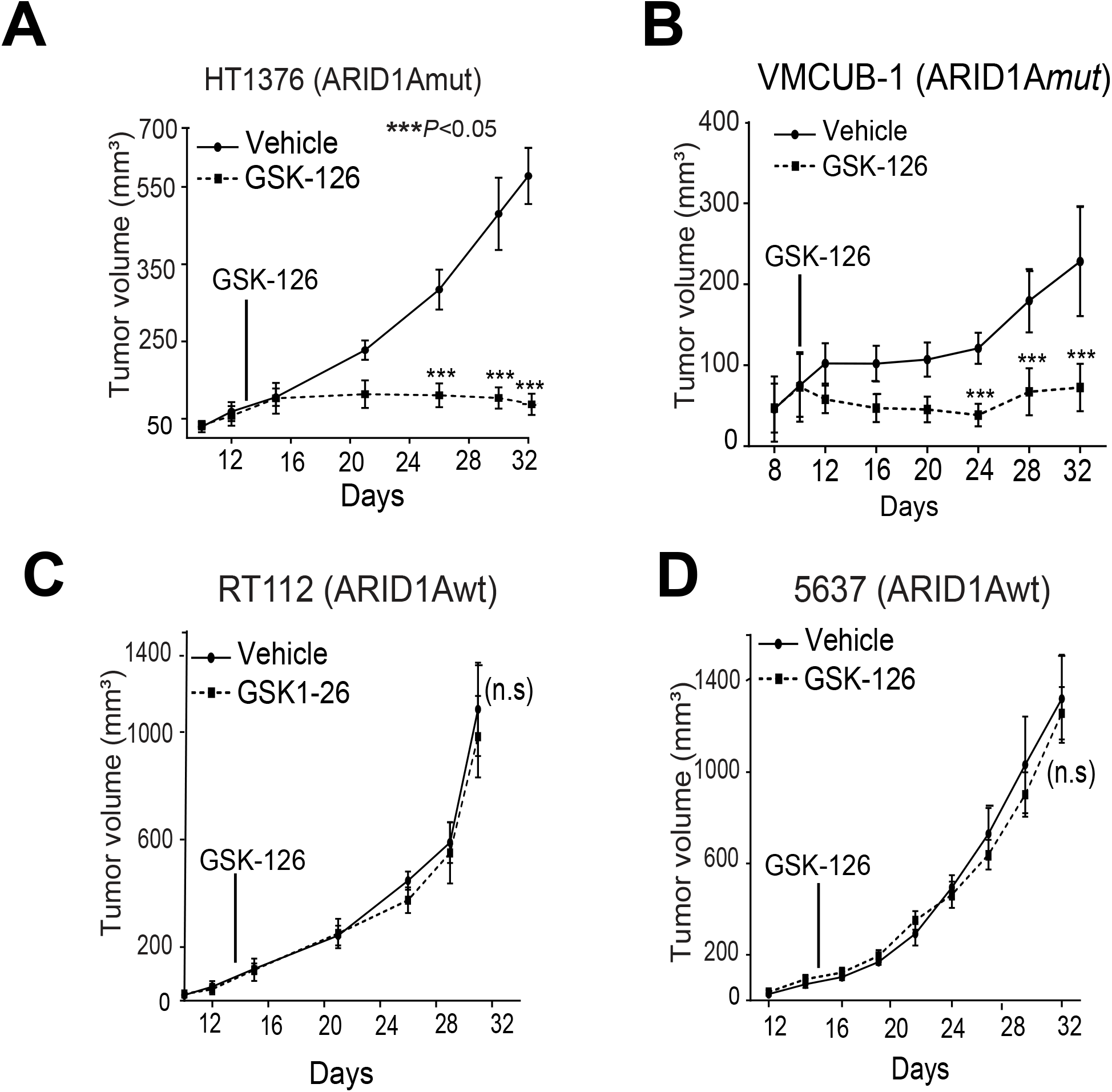
Bladder cancer xenografts harboring ARID1A mutations are sensitive to EZH2 inhibition. (A-D) Plots of tumor volume at indicated days after mice inoculated with HT1376 (ARID1A mutant), VMCUB-1 (ARID1A mutant), RT-112 (ARID1A wild type), and 5637 (ARID1A wild type) cells respectively, treated with EZH2 inhibitor GSK126 (dashed line) or vehicle (solid line). “ns” – non-significant.

### ARID1A knockdown induces GSK-126 sensitivity in ARID1A wild-type bladder cancer cells

To investigate whether ARID1A deficiency is sufficient for EZH2 inhibitor sensitivity, we generated ARID1A-wildtype bladder cell lines harboring stable shRNA-mediated knockdown of ARID1A. Notably, these cell lines showed ARID1A protein levels that are comparable to ARID1A mutant cell lines (**Fig 4A**). While ARID1A knockdown did not affect colony formation or cell viability at baseline, ARID1A knockdown did result in increased sensitivity to GSK-126 as evidenced by decreased colony formation (**Fig 4B**), decreased viability (**Fig 4C**), and increased pharmacologic sensitivity (10-fold decrease in IC50) (**Fig 4D**). These ARID1A stable knockdown cells were then used to generate xenografts to test their sensitivity to EZH2 inhibition *in vivo*. While RT112 and 5637 ARID1A wild-type cells were previously resistant to GSK-126 treatment (**Fig 3 C, D**), ARID1A knockdown cells were quite sensitive and their growth was nearly completely inhibited (**Fig 4 E, F**). Taken together, it was confirmed that sensitivity of bladder cancer cells towards an EZH2 inhibitor is dependent on ARID1A deficiency.

**Figure 4:**
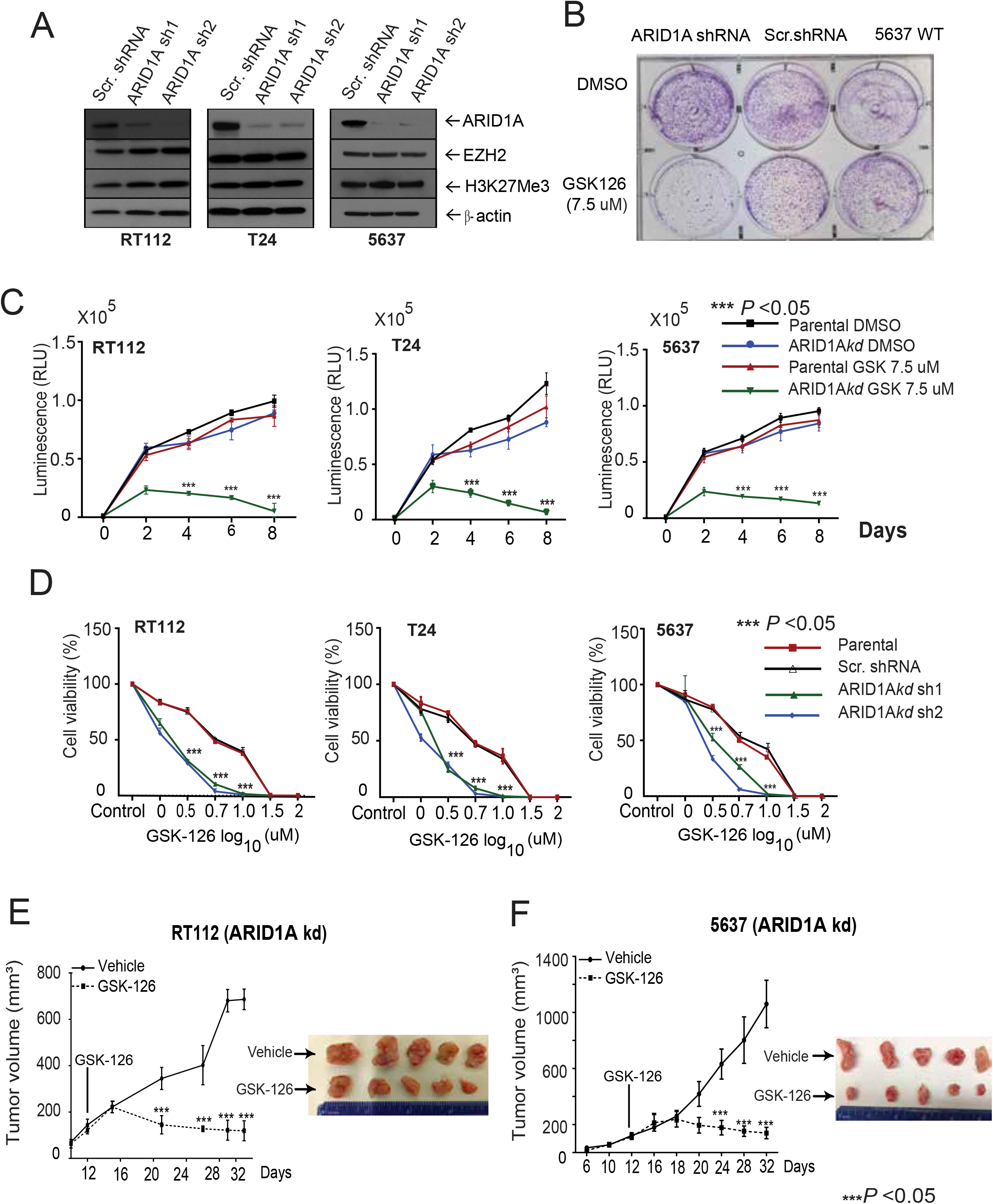
ARID1A knockdown sensitizes bladder cancer cells to EZH2 inhibition. (A) Immunoblots showing expression of ARID1A, EZH2 and tri-methylated H3K27 (H3K27me3) in ARID1A wild type BLCa cell lines after ARID1A stable knockdown (KD) with two separate shRNA sequences along with scrambled (scr) shRNA. (B) Colony formation assays of untreated, scramble shRNA, and ARID1A KD 5637 cells with DMSO or GSK126 treatment. (C) Plots demonstrate the cell proliferation in parental (scr) ARID1Awt and KD cells and (D) Plots show the dose-response viability curves of ARID1A wild type (T24, 5637 and RT112) bladder cancer cell lines with and without ARID1Akd treated with GSK126 for 144h. (E, F) Plots of tumor volume at indicated days after mice inoculated with RT-112 (ARID1A knockdown), and 5637 (ARID1A knockdown) cells respectively, treated with EZH2 inhibitor GSK126 (dashed line) or vehicle (solid line). Inset, photomicrographs of xenograft tumors.

## Discussion

Metastatic urothelial carcinoma is generally incurable with modest survival benefit provided by cisplatin-based first-line chemotherapy (median survival ∼15 months) (24-31). Durable benefits with post-platinum PD-1/L1 inhibitors extend to a minority of patients (∼20%) and the median survival is <1 year (25, 32-36)(37, 38). Third-line salvage therapies provide incremental benefits with median overall survival of ∼1 year and are not curative (e.g. Enfortumab Vedotin). Moreover, the first targeted agent, erdafitinib, was recently shown to be active and approved to treat post-platinum patients with activating somatic FGFR2/3 mutations or fusions (39-43). Hence, new therapeutic approaches are critically needed to yield cures, which will only arrive with better understanding of mechanisms of resistance and therapeutically actionable targets. Given the heterogeneity of this malignancy with multiple genomic alterations, there remains a role for rational approaches targeting these subsets of patients.

Recent studies have revealed that ARID1A is frequently mutated across a wide variety of human cancers including bladder, gastric, pancreatic, and ovarian cancers, and also has *bona fide* tumor suppressor properties (6, 7). As ARID1A is the DNA-binding subunit of the large ∼1.15MDa SWI/SNF multi-subunit complex, its loss through non-sense and missense mutations is thought to result in complex disassembly. Interestingly, loss of just one allele of ARID1A results in embryonic lethality in mice (44). Clearly, the protein levels of ARID1A are important in development and disease.

While loss of tumor suppressors like ARID1A can be difficult to therapeutically target directly, oftentimes these losses result in therapeutic vulnerabilities that can be targeted through a synthetic lethality approach. Using various bladder cancer cell lines with ARID1A truncating mutations or shRNA-mediated depletion, our experiments reveal that EZH2 inhibition is synthetically lethal in bladder cancer cells with ARID1A deficiency. Notably, similar findings have been discovered in ovarian clear cell carcinoma (15). Other groups have independently investigated this relationship in bladder cancer cells and come to somewhat different conclusions (23). On careful review, the cells in these studies were only treated with EZH2 inhibitor for 2-3 days and showed no specific sensitivity for EZH2 inhibition, whereas we have found that at least 6-8 days of treatment is necessary to see maximal differences in viability. This alone could explain the differences between our results.

Pharmacologic inhibitors of EZH2 are currently being investigated in a large variety of tumor types including lymphoma, sarcomas, and advanced treatment resistant solid tumors (reviewed in ref (45)). B-cell lymphomas often harbor activating mutations in EZH2, and some sub-types of sarcomas harbor mutations in SWI-SNF subunits SMARCB1 or SMARCA4. In fact, the EZH2 inhibitor tazemetostat was approved by the FDA in 2020 for the treatment of advanced epithelioid sarcoma. Thus, there is strong rationale to re-purpose EZH2 inhibitors for the pharmacologic treatment of bladder cancer patients whose tumors harbor ARID1A-mutations and/or deficiency. Notably, there is currently an active phase I/II trial investigating combination pembrolizumab and the EZH2 inhibitor tazemetostat in molecularly unselected advanced urothelial carcinoma (clinicaltrials.gov NCT03854474). The results herein suggest that sub-group analyses of this and other trials should focus on patients with ARID1A-deficient tumors.

We are currently actively investigating the molecular mechanisms of how ARID1A loss leads to EZH2 inhibitor sensitivity in bladder cancer. In ovarian carcinoma with clear cell histology, ARID1A and EZH2 both compete (along with histone deacetylases) to modulate the transcriptional activity of the PIK3IP1 gene (15, 46). PIK3IP1 is a 47kDa transmembrane protein which has been shown to directly bind to and inhibit the catalytic subunits of Class I PI3K, including PIK3CA (aka p110α) (47, 48). Thus, these studies showed that ARID1A loss combined with EZH2 inhibition results in a de-silencing of PIK3IP1 which inhibits PI3K/AKT/mTOR signaling resulting in inhibition of cellular proliferation. The PI3K/AKT/mTOR pathway is important in bladder cancer tumorigenesis, as a significant proportion of these tumors harbor activating mutations in PIK3CA (2). We therefore hypothesize that ARID1A loss in bladder cancer may result in PI3K/AKT/mTOR pathway activation and dependence that can be indirectly targeted by EZH2 inhibitor-mediated upregulation of PIK3IP1. Further studies are underway to investigate this hypothesis. In summary, our studies suggest a rationale for treating ARID1A mutated bladder cancers by targeting EZH2 with specific small molecule inhibitors.

