## Supplemental figure legends for "ARID1A-mutant and deficient bladder cancer is sensitive to EZH2 pharmacologic inhibition"

**Supplementary Figure Legends**:

**Supplementary Figure 1**: Photomicrographs of xenograft tumors from mice inoculated with (A) RT-112 (ARID1A wildtype), (B) 5637 (ARID1A wildtype) and (C) HT1376 (ARID1A mutant) cells.

**Supplementary Figure 2**: Western blot showing the expression of tri-methylated H3K27 (H3K27me3) in flank tumor bearing (nu/nu) mice treated with GSK126 (100 mg/kg/once/daily) for 21 days in (A) HT1376 (ARID1A mutant), (B) RT112 (ARID1A wildtype) and (C) RT112 (ARID1A knockdown) cell lines. Treatment with GSK126 for 21d was started after the tumor volume reached to 150 to 200 mm^3^ in size with vehicle (Captisol).
