## Supplementary figures and images for "ARID1A-mutant and deficient bladder cancer is sensitive to EZH2 pharmacologic inhibition"

### Supplemental Figures

#### Supplementary Figure 1

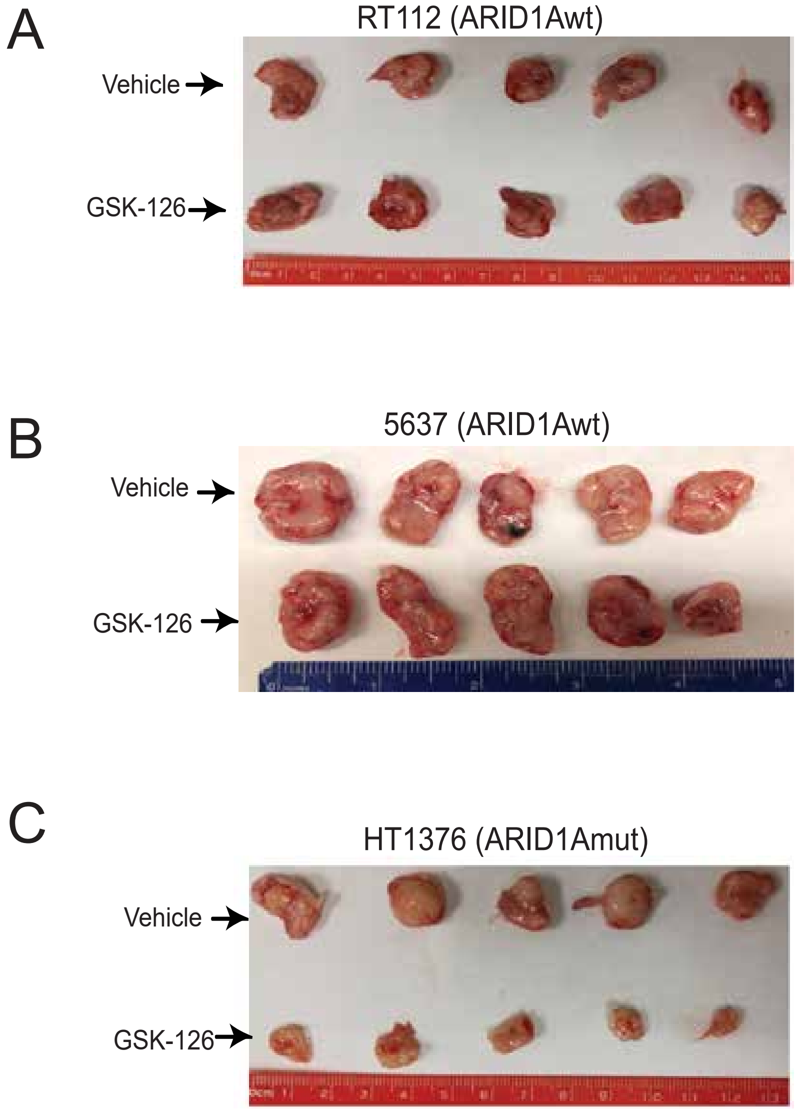

#### Supplementary Figure 2

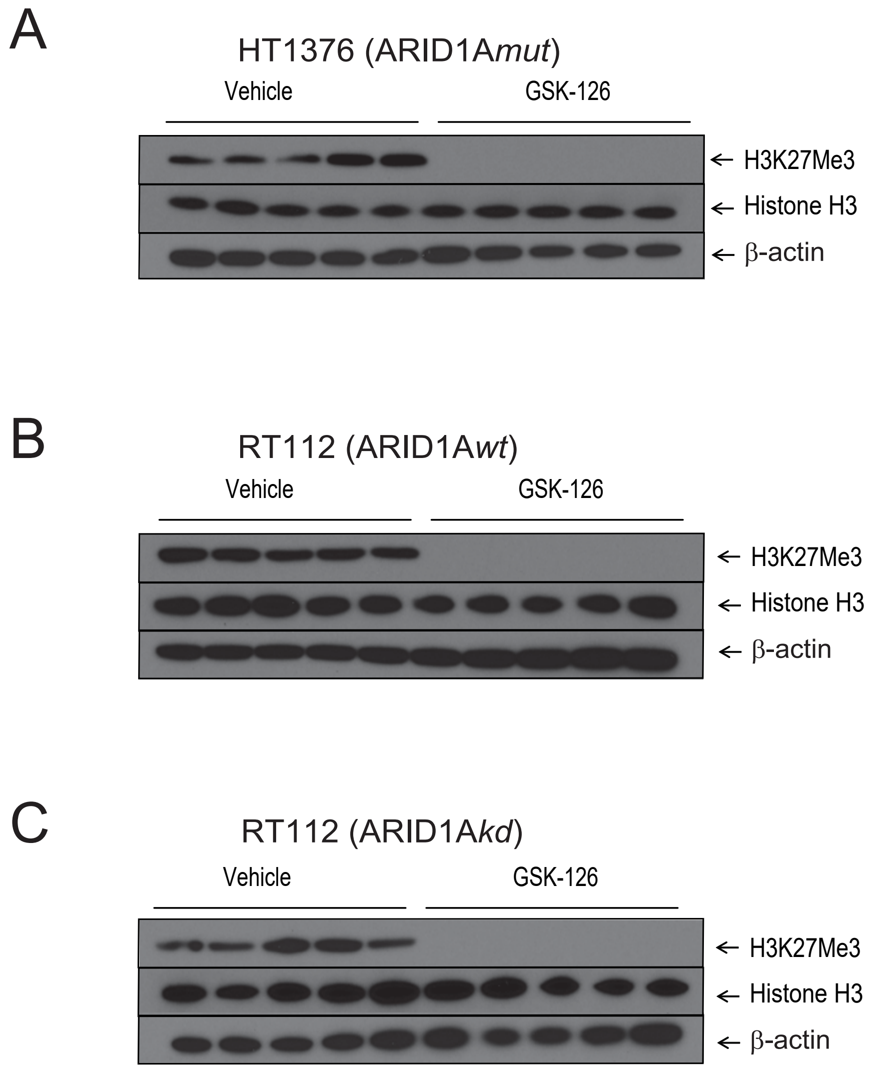
